# Mesophyll-specific overexpressing of *SvHXK6* gene improves water use efficiency without yield loss in C_4_ *Setaria viridis*

**DOI:** 10.1101/2023.07.10.548403

**Authors:** Yogesh Chaudhari, Lily Chen, Shahasad Salam, Matthew Paul, Robert Furbank, Oula Ghannoum

## Abstract

Hexokinases (HXK) were the first sugar signalling proteins identified in plants and are well known for their feedback regulation of photosynthetic gene expression. In some C_3_ plants, HXKs have been found to regulate stomatal function. However, the role of HXK in C_4_ photosynthesis, which is inherently more water use efficient than C_3_ metabolism, remains poorly understood. Here, we report on the first tissue-specific modification of HXK in a C_4_ plant. *SvHXK6* was expressed in the model C_4_ grass *Setaria viridis* under the control of the *ZmPEPC* promoter (*ZmPEPC*_*pro*_), which directs expression in the leaf mesophyll tissue. Three *S. viridis* transgenic lines with increased abundance of *SvHXK6* transcripts in the leaf tissue showed significant reduction in stomatal conductance with minimal effects on leaf CO_2_ assimilation rate. Consequently, the transgenic lines had higher leaf-level water use efficiency relative to the control (wild-type and null) plants. Overexpression of *SvHXK6* had no effect on shoot biomass or seed yield of the *S. viridis* plants. Our study shows conserved function of HXK in regulating stomatal conductance in a C_4_ grass, demonstrating possible widespread utility in improving water use efficiency in C_4_ as well as C_3_ species.

## Introduction

More creative ways are required to improve crop performance, including of C_4_ crops, in the face of increasingly variable rainfall and warmer temperatures. Traits related to water use are critical for crop productivity and survival. Whole plant WUE (*WUE*_*P*_), the ratio of biomass produced to water used, is an important determinant of crop yield, especially under water limitation (Leakey et al., 2019). At the leaf level, intrinsic water use efficiency (*iWUE*) is defined as the ratio of net CO_2_ assimilation rate (*A*_*n*_) to stomatal conductance to water vapour (*g*_*s*_). Greater *WUE* may potentially lead to greater crop yield if it results from increased *A*_*n*_ (biomass accumulation) rather than reduced g_s_ (water loss) (Farquhar et al., 1989).

C_4_ photosynthesis saturates at relatively low CO_2_ concentration due to the operation of a CO_2_ concentrating mechanism (CCM) and the high affinity of PEP carboxylase for CO_2_ (Hatch, 1987). Consequently, C_4_ plants tend to operate at lower *g*_*s*_ and achieve higher *iWUE* relative to C_3_ plants (Ghannoum et al., 2011). However, global climate change and increasingly variable rainfall patterns are impacting food production in tropical and subtropical regions where C_4_ crops are grown (Watson-Lazowski and Ghannoum, 2021). Hence, for securing future food production in these regions it is critical to improve *iWUE* of C_4_ crops, which requires a better understanding of the mechanisms regulating *iWUE* in these plants. Owing to the CCM and the near CO_2_-saturation of C_4_ photosynthesis, reduced g_s_ is expected to have a smaller impact on *A*_*n*_ in C_4_ relative to C_3_ photosynthesis (Ghannoum et al., 2011). Indeed, *iWUE* is generally better correlated with g_s_ than *A*_*n*_ in C_4_ plants (Cano et al., 2019). Therefore, a targeted strategy which reduces *g*_*s*_ may be particularly beneficial for C_4_ crops by slightly boosting their *iWUE* without compromising *A*_*n*_.

Hexokinases (HXKs) are essential metabolic catalysts which also act as glucose sensors that can repress the expression of some photosynthetic genes in response to high internal glucose concentrations (Moore et al., 2003). More recently, the specific expression of *AtHXK* in guard cells of C_3_ dicots, such as tobacco, reduced *g*_*s*_ and enhanced *WUE* (Lugassi et al., 2019). Evidence indicated that HXK senses the accumulation of sugars in the apoplast, leading to the efflux of water out of the guard cells and hence, stomatal closure (Kelly et al., 2013). It seems that HXK promotes the production of nitric oxide (NO) and hydrogen peroxide (H_2_O_2_) in guard cells (Shen et al., 2021), which interacts with the signalling pathways of plant hormone abscisic acid (ABA), leading to stomatal closure (Kelly et al., 2013). However, the role of HXKs in regulating stomatal conductance of C_4_ plants, which are inherently more water use efficient than C_3_ plants, remains poorly understood.

Genetic manipulation of sensing *HXKs* in C_3_ plants has been mostly carried out using the constitutive cauliflower mosaic virus *35S* promoter for whole plant manipulation (Dai et al., 1999). More recently, a guard cell specific promoter was successfully used in C_3_ dicot plants (Kelly et al., 2013). With the exception of manipulating *HXK* in maize mesophyll protoplasts, no genetic modification of HXKs has been carried out in any C_4_ species, and none using cell type-specific promoters. Additionally, a guard cell promoter has yet to be successfully validated in C_4_ monocots (Kelly et al., 2017).

To address our lack of knowledge on the role of HXKs in C_4_ photosynthesis and stomatal function, we overexpressed *SvHXK6* (homologue of sensing *AtHXK1*) in the model C_4_ monocot *Setaria viridis* under the control of the *ZmPEPC* promoter (*ZmPEPC*_*pro*_), which directs expression in the leaf mesophyll tissue (Sattarzadeh et al., 2010). Our aim was to specifically study the function of this putative sugar sensor within the source leaf of a C_4_ plant, without pleiotropic effects on other plant processes. Three stable *S. viridis* transgenic lines with significantly higher abundance of *SvHXK6* transcripts were obtained and analysed.

Based on previous studies in C_3_ plants, overexpression of sensing *HXK* in the leaf tissue is expected to reduce photosynthesis and plant growth as a result of heightened feedback inhibition of photosynthetic genes expression (Moore et al., 2003). In addition, overexpression sensing *HXK* is expected to reduce stomatal conductance and increase *iWUE* (Kelly et al., 2013). However, we recently showed that C_4_ photosynthesis is little affected by increased glucose concentration following shifts to hight light in the C_4_ model grass *S. viridis* (Henry et al., 2020). Therefore, we expected that the overexpression of *SvHXK6* under the control of *ZmPEPC*_*pro*_ in *S. viridis* will mostly illicit a stomatal closure response rather than a photosynthetic feedback inhibition response. Consequently, we expected to see improvements in *iWUE* without reductions in productivity.

## Materials and Methods

### Transformation of Setaria viridis

The sugar sensing gene *SvHXK6* in *Setaria viridis* was identified based on maize *ZmHXK6* and rice *OsHXK6* sequences in NCBI (**Figure S1**). *S. viridis* lines overexpressing the sugar signalling gene *SvHXK6* were generated using the maize phosphoenolpyruvate carboxylase promoter (*ZmPEPC*_*pro*_) for the preferential expression in mesophyll cells (Wang et al., 2017), which was confirmed using the Bio-Analytical Resources (BAR) for Plant Biology (**Figure S2**). Stable transformation of *S. viridis* (accession ME034V) via *Agrobacterium tumefaciens* strain AGL1 and Gateway® cloning (**Figure S3**) was carried out as originally described in (Van Eck, 2018), using a modified protocol from (Osborn et al., 2017).

### Verification of transformants and homozygosity

Genomic DNA was extracted from leaves of putative T_0_ transformant plants and genomic PCR was performed for the detection of gene encoding selectable marker *hygromycin phosphotransferase* (*hpt)*. Droplet digital PCR was performed to determine the number of transgene copies. The single copy T_0_ plant lines were selected, the seeds were harvested and further grown to T_1_ generation. T_1_ lines that doubled in transgene copy number were deemed homozygous (**Tables S1 and S2**). RNA was extracted from all homozygous lines to quantify the *SvHXK6* transcripts from all *S. viridis* homozygous *HXK6* T_2_ mutants using semi-quantitative PCR to get an initial indication of overexpression, which was verified for three *OE* lines using RT-qPCR (**Figure S4**).

### Plant Culture and Growth analysis

*Setaria viridis* (ecotype ME034V) seeds (wild type, null and transformants) were germinated and grown as described in (Henry et al., 2020) in controlled environmental chambers (CONVIRON PGR 15) under the following conditions: light intensity of 500 μmol m^-2^ s^-1^; 16h day: 8h night, 28/22°C day/night; 400 ppm CO_2_ and 60% relative humidity. Three harvests (n=4) were performed at 10 days (seedlings), 20 days (vegetative), and 30 days (∼8-9 days after panicles had emerged) after germination. At harvest, plant height was measured then leaves, stems and panicles were separated and dried at 70°C before weighing. Leaf mass per area (LMA) was measured on last fully expanded leaves.

### Leaf gas exchange measurements and soluble sugars

Gas exchange measurements (n=4) were carried out using a Li-COR 6400XT portable photosynthesis system after 15 days of germination on the youngest fully expanded leaf from the main stem. The response of CO_2_ assimilation rate to step increases of intracellular CO_2_ (*A-C*_*i*_ curve) was measured at a standard light intensity of 1000 μmol m ^-2^ s ^-1^ around midday. Leaf discs were collected around midday from leaves used for gas exchange, and snap frozen in liquid nitrogen. Samples were ground using a TissueLyser II (QIAGEN), and extracted 3-times in 80% methanol at 90°C for 30 min. The extract was concentrated using a vacuum evaporator (EPPENDORF, Hamburg, Germany), and resuspended in MilliQ water. Soluble sugars were analysed using a kit (MEGAZYME, Bray, Ireland).

### Statistical analysis

To assess the statistical differences between WT, nulls and transgenic lines, one-way ANOVA was carried out using R software (R Core team, 2021). If there was a significant difference (p ≤ 0.05), an Agricolae function was used to rank the means. Regression analysis was carried out in R using linear modelling (lm).

## Results

### Confirmation of SvHXK6 overexpression by RT qPCR

The *SvHXK6* RNA was quantified from homozygous lines which were previously verified using digital droplet genomic PCR. Using semi-quantitative PCR, the *SvHXK6* gene was compared to two housekeeping genes (*ACTIN* and *GAPDH*). Three homozygous lines (*SvHXK6-30 OE* (H30), *SvHXK6-43 OE* (H43) and *SvHXK6-44 OE* (H44)) showed higher (∼2-fold) levels of *SvHXK6* expression relative to the WT (**Figure S4A**). These results were confirmed by qPCR, but with a lower level of overexpression (∼1.4-fold) relative to the WT (**Figure S4B**). Only one line (H43) showed significantly higher overexpression. Consequently, three lines, *SvHXK6-30 OE, SvHXK6-43 OE* and *SvHXK6-44 OE*, were chosen for further analysis. A line which underwent transformation but had no transgene in T_0_ plants was treated as a *null* mutant and used as a negative control (**Figure S5**).

### Leaf gas exchange and soluble sugars

Under standard conditions, both WT and null plants showed similar CO_2_ assimilation rate (A_net_), stomatal conductance (g_s_) and intrinsic water use efficiency (*iWUE*) (**Figure 1A-C, Table 1**). A_net_ was significantly lower in the *SvHXK6-30 OE* mutant relative to the other lines, while g_s_ was lower in all three *OE* lines relative to control (WT and null) plants (**Figure 1A-B**). *iWUE* tended to be higher in all three *OE* lines relative to control plants, and this increase was significant in the H30 plants (**Figure 1C, Table 1**). Across all measurements, *iWUE* showed strong negative correlations with C_i_ (r^2^=0.99, *p*<0.001), g_s_ (r^2^=0.88, *p*<0.001), and weakly with A_net_ (r^2^=0.23, *p*>0.05) (**Figure 1D-F**).

**Table 1.** **Summary of averages** of phenotypic data for *S. viridis* WT, null and *SvHXK6 OE* transgenic lines, and their statistical differences. The significance was calculated using one-way ANOVA. Values in bold red indicate significant difference with both WT and null plants.

|  |  |  | *S. viridis* transgenic lines<br>overexpressing *SvHXX6* |  |  |
| --- | --- | --- | --- | --- | --- |
| Parameters | WT | Null | H30 | H43 | H44 |
| **Harvest 1** |  |  |  |  |  |
| Plant height (cm) | 19.07 a | 18.15 a | 18.3 a | 18.1 a | 18.2 a |
| Shoot dry mass (g) | 0.53 a | 0.55 a | 0.50 a | 0.55 a | 0.62 a |
| **Harvest 2** |  |  |  |  |  |
| CO<sub>2</sub> assimilation rate ($\mu\text{mol}^{-2} \text{s}^{-1}$) | 23.09 a | 23.8 a | **18.7 b** | 21.1ab | 21.0 ab |
| Stomatal conductance ($\text{mol m}^{-2} \text{s}^{-1}$) | 0.24 a | 0.24 a | **0.14 b** | **0.16 b** | **0.16 b** |
| *iWUE* ($\mu\text{mol CO}_2 (\text{mol H}_2\text{O})^{-1}$) | 100 a | 96 a | **134 b** | 128 ab | 127 ab |
| Initial slope of *A-Ci* curves | 0.21 a | 0.19 a | 0.20 a | 0.21 a | 0.19 a |
| *A*<sub>max</sub> of *A-Ci* curves ($\mu\text{mol}^{-2} \text{s}^{-1}$) | 23.36 a | 24.6 a | **19.4 b** | 22.7 ab | 21.8 ab |
| Sucrose (mg/g FW) | 10.01 a | 0.86 a | 0.97 a | 0.79 a | 0.84 a |
| Glucose (mg/g FW) | 0.58 ab | 0.042 ab | 0.074 ab | 0.082 a | 0.036 b |
| Fructose (mg/g FW) | 0.51 a | 0.047 a | 0.051 a | 0.059 a | 0.056 a |
| LMA ($\text{g m}^{-2}$) | 42 a | 42 a | 48 a | 41 a | 42 a |
| Plant height (cm) | 43.4 ab | 39.8 bc | 38.7 c | 39.1 c | 44.2 a |
| Shoot dry mass (g) | 1.97 bc | 1.70 c | 2.33 ab | 2.28 abc | 2.59a |
| **Harvest 3** |  |  |  |  |  |
| Plant height (cm) | 120 a | 116 b | 115 b | 116 b | 116 b |
| Shoot dry mass (g) | 22.7 a | 21.6 a | 22.0 a | 21.9 a | 21.17 a |
| Panicle dry mass (g) | 3.55 a | 3.41 a | 3.21 a | 3.34 a | 3.15 a |
| Leaf fraction (% of shoot biomass) | 26.8 | 24.4 | 32.1 | 29.9 | 28 |
| Stem fraction (% of shoot biomass) | 57.6 | 59.8 | 53.3 | 54.8 | 57.2 |
| Panicle fraction (% of shoot biomass) | 15.6 | 15.8 | 14.5 | 15.3 | 14.8 |
| Relative growth rate ($\text{g g}^{-1} \text{d}^{-1}$) | 0.081a | 0.080ab | 0.082a | 0.080ab | 0.076b |
| Shoot elongation rate (cm cm<sup>-1</sup> d<sup>-1</sup>) | 0.040a | 0.040a | 0.040a | 0.040a | 0.040a |

**Figure 1.**
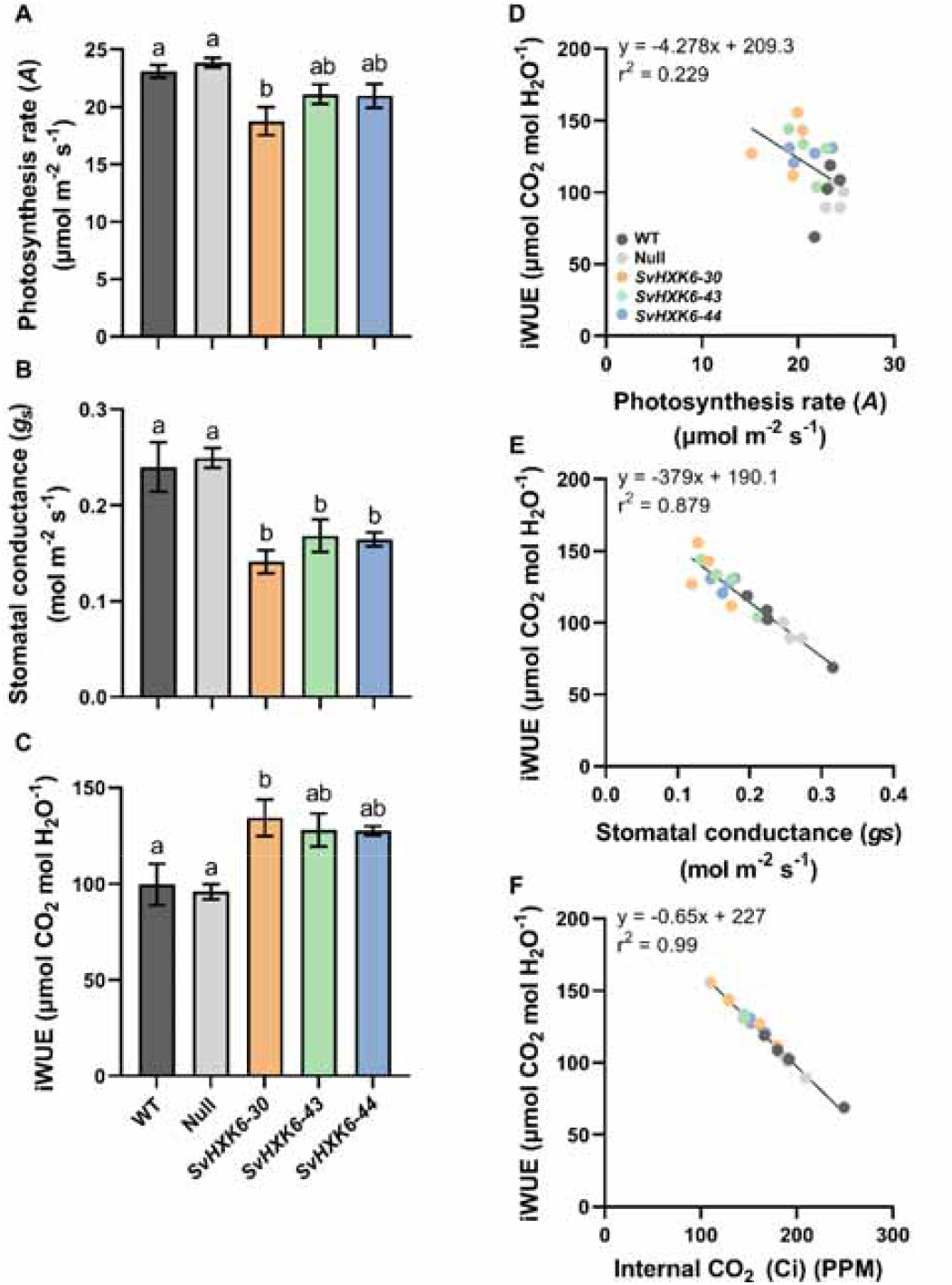
Leaf gas exchange measurements of 20 days old *S. viridis* WT, null and *SvHXK6 OE* mutants. CO_2_ assimilation rate, A_net_ (A), (B) stomatal conductance, g_s_, (C) intrinsic water use efficiency (*iWUE* = A_net_/g_s_), and *iWUE* as a function of (D) A_net_, (E) g_s_, and (F) intercellular CO_2_ concentration (C_i_) of WT (dark grey), null (light grey), H30 (orange) H43 (green) and H44 (blue) plants (n=4). All measurements were made using the fully expanded source leaves at mid-day using a LI-6400XT at 26 °C leaf temperature at an irradiance of 1000 μmol photons m^-2^ s^-1^. Error bars represent standard errors. Different letters above the bars denote the significant differences and same letters indicate similarity among the WT, null and transgenic plants. Significance is calculated using one-way ANOVA.

Analysis of the response of *A-C*_*i*_ curves (**Figure 2A**) revealed no differences in the initial slope between the *S. viridis* lines (**Figure 2B**). Maximal or CO_2_-saturated photosynthesis rate (A_max_) was significantly lower in only the *SvHXK6-30 OE* mutant relative to the other lines (**Figure 2C**), confirming the trends observed with A_net_ (**Figure 2A**).

**Figure 2.**
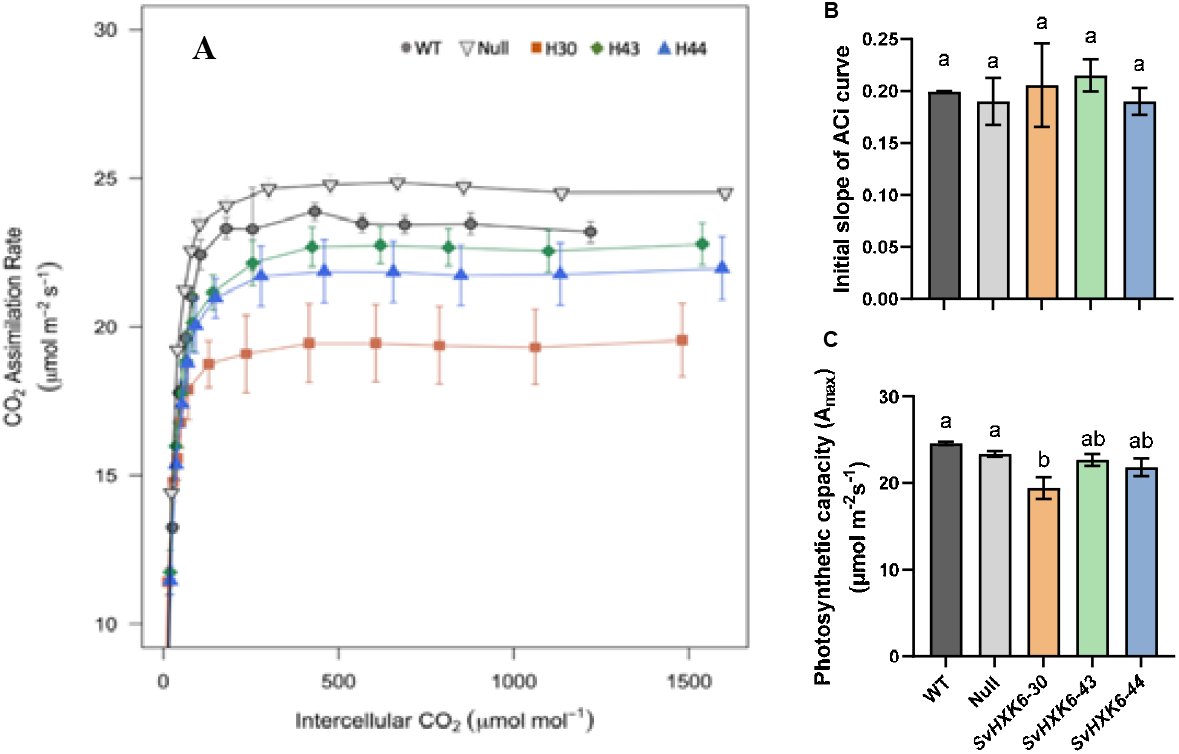
Characteristics of the *A-C*_*i*_ curves of 20 days old *S. viridis* WT, null and *SvHXK6 OE* mutants. (A) The response of net assimilation to internal CO_2_ concentration (*A-C*_*i*_ curve), (B) initial slope and (C) maximal CO_2_ saturated assimilation rate (A_max_) of the *A-C*_*i*_ curves of WT (dark grey), null (light grey), H30 (orange), H43 (green) and H44 (blue) plants (n=4). All measurements were made using the fully expanded leaves of the main stems at mid-day using a LI-6400XT at 26°C leaf temperature at an irradiance of 1000 μmol photons m^-2^ s^-1^. The *A-Ci* curves were measured at the following CO_2_ concentration (50, 100, 150, 200, 250, 300, 400, 600, 800, 1000, 1200, 1500, and 2000 ppm). Before starting the curve, the leaf was allowed to stabilize for 10-15 min, and an initial measurement was recorded at 400 CO_2_ ppm. Error bars indicate standard error of the mean (SEM). Different letters above the bars denote the significant differences and same letters indicate similarity among the WT, null and transgenic plants. Significance is calculated using one-way ANOVA.

Sucrose was the major soluble carbohydrate present in the source leaves while glucose and fructose levels were considerably lower in all *S. viridis* plants. No significant differences were observed in any of the soluble sugars between the transgenic and the control plants (**Figure 3, Table 1**).

**Figure 3.**
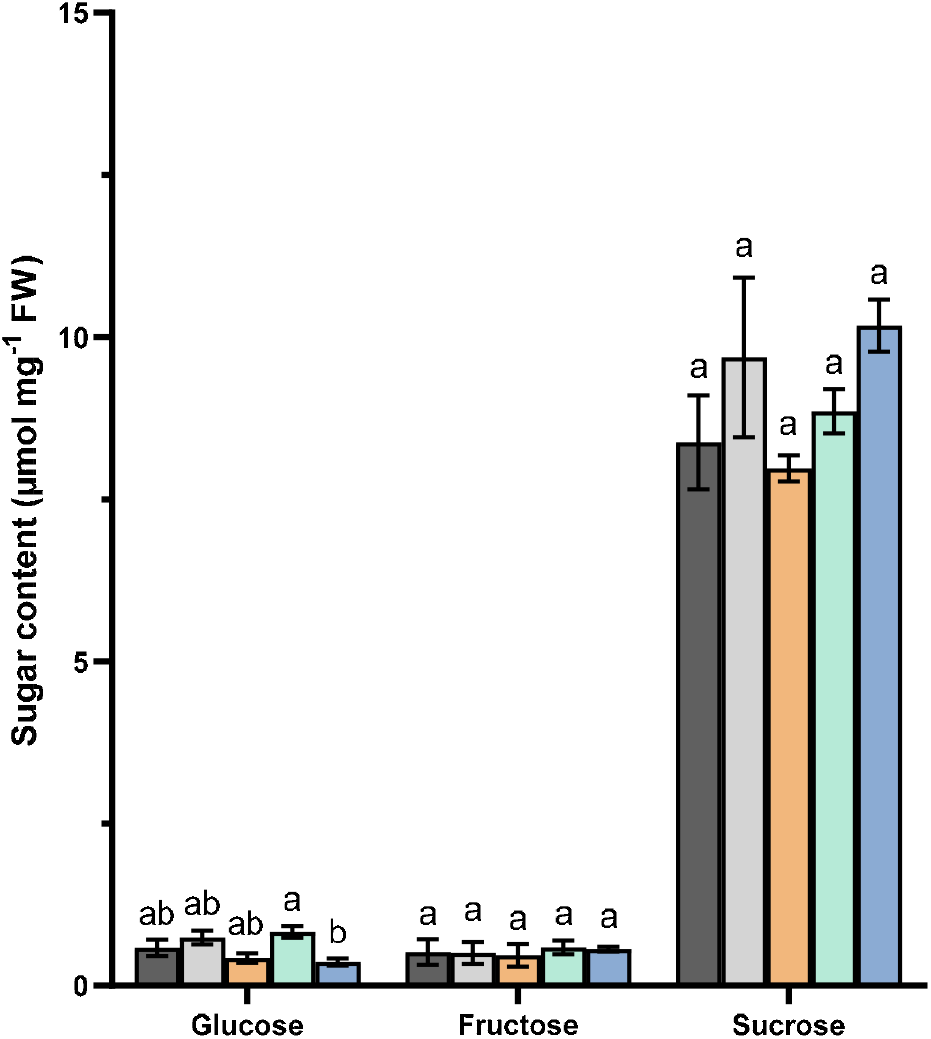
Sugar contents in the source leaves of *S. viridis SvHXK6 OE* mutants. Glucose, fructose and sucrose were extracted from leaf samples harvested at midday from 20 days old WT (dark grey), null (light grey), H30 (orange), H43 (green) and H44 (blue) plants (n=4). Measurements were taken using spectrophotometer and related to standard curve based on glucose detection. Error bars represent standard errors. Different letters above the bars denote the significant differences at the 5% level, and same letters indicate similarity among the WT, null and transgenic plants. Significance is calculated using one-way ANOVA.

### Plant growth and biomass allocation

There were no significant differences between the transgenic and control plants, or between the WT and null plants in any of the measured growth parameters, including LMA, plant height, shoot biomass, relative shoot elongation rate, SER (**Table 1**), relative growth rate, RGR (**Figure 4A**), and yield, calculated as the panicle weight (**Figure 4B**). The proportion of biomass allocation to leaves was slightly higher (∼2.5% more biomass allocated to leaves) in the three *OE* lines relative to the control plants. The biomass fraction allocated to the panicles remained similar across the *S. viridis* lines (**Figure 4C, Table 1**).

**Figure 4.**
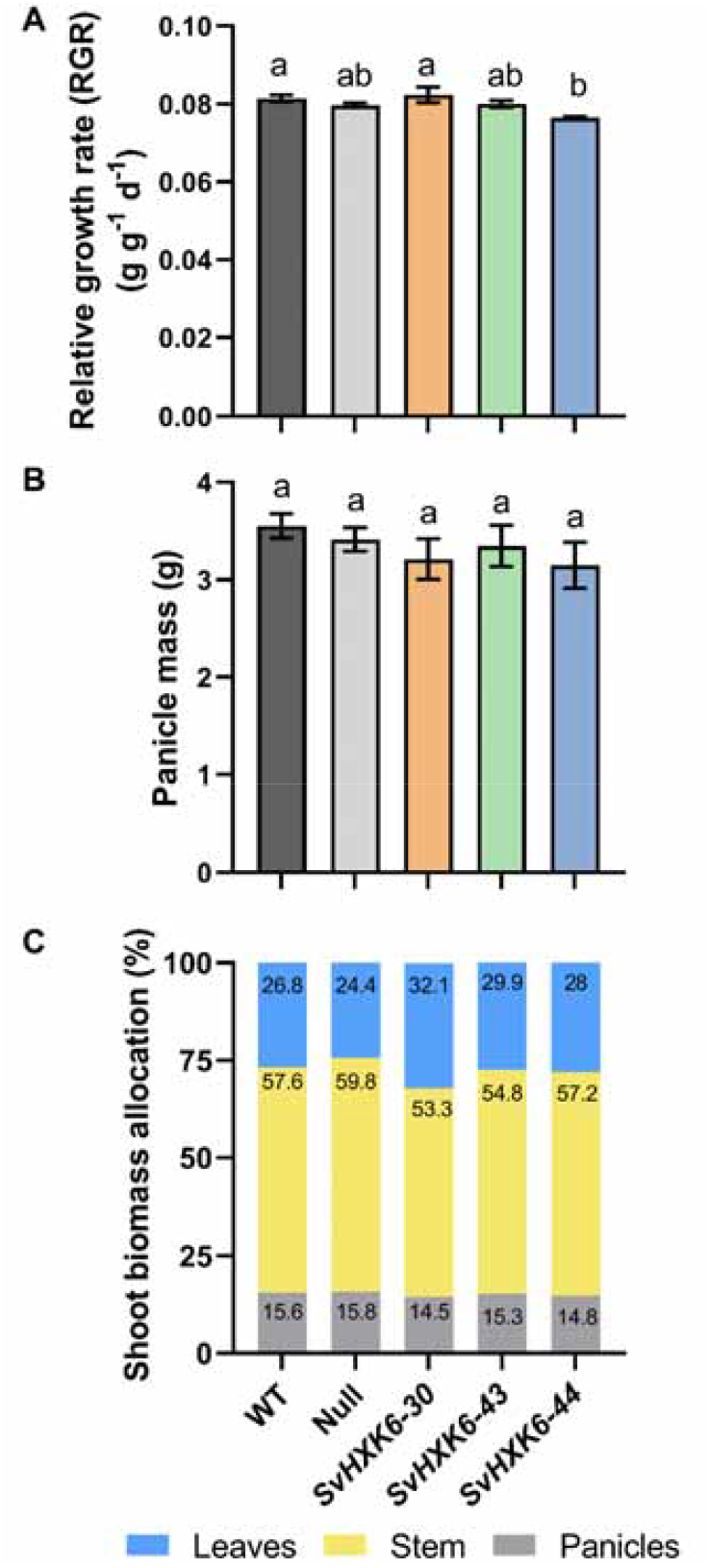
Plant growth characteristics of *S. viridis* WT, null and *SvHXK6 OE* mutants. (A) relative growth rate of the shoot (RGR), (B) panicle dry mass, and (C) biomass allocation to leaves, stems, and panicles of WT (dark grey), null (light grey), H30 (orange), H43 (green) and H44 (blue) plants (n=4). Plants were harvested at 10, 20 and 30 days after germination. Shoot biomass consisted of leaves, stems, and panicles. RGR was calculated as the linear slope of the natural logarithm of dry weight over time for the shoot biomass. Panicle dry mass and biomass allocation were measured on 30 days old pants. Different letters above the bars denote the significant differences at

## Discussion

### Conserved function of sensing HXKs between C_3_ and C_4_ plants

We overexpressed *SvHXK6*, a homologue of sensing *AtHXK1* (**Figure S1**), in the mesophyll tissues of the C_4_ grass *S. viridis* under the control of *ZmPEPC*_*pro*_. Relative to the control (WT and null) plants, only one *OE SvHXK6* line showed significant reduction in the rate of photosynthesis, while stomatal conductance was significantly reduced in all three transgenic lines (**Figure 1**). No effect was observed on plant productivity or yield (**Figure 4**).

Overexpression of sensing *AtHXK1* in the C_3_ plants Arabidopsis and tomato plants under the control of the constitutive *35S* promoter reduced the expression of photosynthetic genes and inhibited plant growth (Dai et al., 1999). In our study, the overexpression of *SvHXK6* may not have been strong enough to heighten feedback inhibition of photosynthetic gene expression, and hence photosynthesis in transgenic *S. viridis* lines. Moreover, we did not observe appreciable differences in soluble sugar content between the control and transgenic *S. viridis* lines (**Figure 3**). The lack of evidence here for sugar build up in the *SvHXK6 OE* lines may be due to the compartmentalisation of sugars between the MC and BSC as well as the fast export of sugars (Lunn and Furbank, 1999; Furbank and Kelly, 2021). Hence, there was not enough sugar accumulation to interact with the slightly increased HXK activity. Recently, we showed that leaf sugar accumulation under a daytime PAR of about 1000 μmol m^-2^ s^-1^ was not high enough to trigger reductions in photosynthesis rate or changes in photosynthetic gene expression in WT *S. viridis* plants grown under non-limiting water and nutrient supplies (Henry et al., 2020). This suggests that C_4_ sugar sensing mechanisms may be inherently less sensitive in comparison to C_3_ plants.

In contrast to constitutive overexpression, the overexpression of *AtHXK1* under the guard-cell specific *KST1* promoter reduced the stomatal conductance and transpiration in both tomato and Arabidopsis plants, without inhibiting plant growth (Kelly et al., 2013). The same was found for citrus plants (Lugassi et al., 2015). These results suggest that HXK plays a specific role in guard cells, influencing stomatal closure (Kelly et al., 2013). In our study, it is likely that the *ZmPEPC*_*pro*_ used to overexpress *SvHXK6* in leaf mesophyll cells was “leaky” and increased HXK activity in guard cells (**Figure S2**). PEPC is well known to be expressed and play an important role in guard cell metabolism (Outlaw, 1990). High level expression of a suite of C_4_ enzymes in guard cells of *Gynandropsis* has been reported from cell specific RNA-seq, including the C_4_ isoform of PEPC (Aubry et al., 2016). Stomatal behaviour was severely affected in a PEPC knock out mutant of *Amaranthus* (Cousins et al., 2007). In the absence of a demonstrated guard-cell-specific promoter for Setaria, or an antibody to detect the Setaria HXK protein in guard cells, this hypothesis remains to be tested. However, these results suggest that, as in C_3_ plants, *SvHXK6* plays a role in regulating stomatal conductance in *S. viridis* plants.

### Overexpressing SvHXK6 as a means of improving crop water use efficiency and productivity under water limitation of C_4_ plants

Overexpression of *SvHXK6* in the mesophyll tissues of the C_4_ grass *S. viridis* increased leaf intrinsic water use efficiency (*iWUE*) in the transgenic lines relative to the WT and null control plants. This effect was significant in the H30 line which had also reduced photosynthesis rates (**Figure 2, Table 1**). These results are particularly significant because *SvHXK6 OE* lines grew as tall as WT and null plants at the final harvest, and no significant difference was recorded in shoot biomass or seed yield between the transgenic and control plants (**Figures 4, Table 1**).

Our results align with a recent study that compared transgenic tobacco lines expressing *AtHXK1* constitutively (*35SHXK2* and *35SHXK5*) and a line with guard-cell targeted overexpression of *AtHXK1* (*GCHXK2*). The study reported significantly better *iWUE* in *GCHXK2* tobacco lines relative to WT when grown under field conditions, without negatively impacting CO_2_ assimilation (Acevedo-Siaca et al., 2022). Given C_4_ photosynthesis is mostly CO_2_-saturated at current ambient CO_2_ concentration, a reduction in stomatal conductance (g_s_) will bring a small reduction in photosynthesis rate (A_net_), due to the CCM, resulting in substantial increase in *iWUE* expressed as A_net_/g_s_ (Ghannoum, 2016).

Breeding for higher *iWUE* has been challenging due to its complexity and low heritability. In addition, selecting for higher *iWUE* may inadvertently result in smaller plants (Ghannoum, 2016). Transgenic approaches to improve WUE have yielded mixed results and revealed that genes involved in stomatal development and guard cell movements influence *iWUE*, but often at the expense of plant productivity (Leakey et al., 2019). Hence, there is a case in exploring further how genetic manipulation of signalling HXKs in C_4_ plants can improve leaf, and hence plant WUE, without compromising plant productivity. Targeting WUE with single genes has been challenging in crops with a few exceptions (Nuccio et al., 2015; Argentina first to market with drought-resistant GM wheat, 2021). Hence targeting HXK in mesophyll or guard cells provides a potentially widespread strategy for improving WUE in C_4_ and well as C_3_ crops. Until a specific guard cell promotor is validated for C_4_ monocots, we expect that the maize PEPC promoter (used in our study) will likely express in Setaria guard cells as the C_4_ isoform of PEPC is also found in the guard cells of C_4_ leaves (Aubry et al., 2016). Under future drier and hotter climates, increases in crop WUE may be essential for sustaining crop yield (Passioura and Angus, 2010; Ghannoum, 2016).

## Conclusions

We showed that overexpression of *SvHXK6* in the leaves of *S. viridis* using a *ZmPEPC*_*pro*_ for MC localisation leads to reduced stomatal conductance and improved *iWUE*, as observed in C_3_ species, but in this case without a reduction in assimilation at air levels of CO_2_. The expected phenotype of reduced photosynthesis resulting from *SvHXK6* overexpression based on the HXK being a key component of feedback regulation of photosynthetic gene expression was observed to a limited degree in only one of the transgenic lines of *SvHXK6 OE*. Reduced stomatal conductance in all transgenic lines supports the hypothesis that *SvHXK6* functions in guard cells as a sugar sensor responding to apoplastic sugar concentration. The study shows a potential universal strategy for targeting of hexokinase to improve WUE in C_4_ as well as C_3_ species with potential widespread utility in stabilising crop performance under increasingly variable rainfall patterns.

## Author Contributions

RF, OG, MP and YC conceived the project. YC collected and analysed the data. SS and LC undertook qPCR analysis. YS, RF and OG prepared the manuscript with input from other authors.

## Acknowledgements

We thank Xuequin Wang who undertook *S. viridis* transformation at Professor Furbank’s lab at Australian National University.

## Data Availability

All data supporting the findings of this study are available within the paper and within its supplementary data published online. Reuse of the data is permitted after obtaining permission from the corresponding author.

